# *Helicobacter pylori* FlgV forms a flagellar motor ring structure required for optimal motility

**DOI:** 10.1101/2022.10.24.513468

**Authors:** Jack M. Botting, Shoichi Tachiyama, Katherine H. Gibson, Jun Liu, Vincent J. Starai, Timothy R. Hoover

**Author notes:** Current address: rPeptide, 1050 Barber Creek, Building 300, Suite 103, Watkinsville, Georgia 30677, U.S.A. J.M.B and S.T. contributed equally to the paper and should considered as co-primary authors.

## Abstract

The bacterium *Helicobacter pylori* has a large flagellar motor that generates significantly higher torque than the archetypical *Escherichia coli* motor. To understand how *H. pylori* navigates the viscous environment of the stomach, it is essential to establish how specific motor components contribute to efficient motility. We show here that the protein FlgV, required for motility in *Campylobacter jejuni*, forms a novel ring associated with the MS and C rings in *H. pylori*. Deletion of *flgV* from *H. pylori* B128 or a highly motile variant of *H. pylori* G27 (G27M) resulted in reduced motility in soft agar medium. Based on comparative analyses of *in-situ* flagellar motor structures of *H. pylori* wild-type and Δ*flgV* mutants, the reduced motility of the Δ*flgV* mutants and the location of the FlgV ring suggest it stabilizes interactions between the MS and C rings and/or plays a role in switching the direction of flagellar rotation. Overall, these results identify a novel motor accessory likely adapted to promote flagellar function for bacterial colonization of high-load environments such as the gastric mucosa.

## INTRODUCTION

The human gastric pathogen *Helicobacter pylori*, a member of the phylum Campylobacterota, uses a cluster of polar sheathed flagella for motility, which is required for colonization in animal models for infection (Eaton *et al*, 1992; Ottemann & Lowenthal, 2002). The bacterial flagellum is organized into three basic parts referred to as the basal body, hook, and filament (Berg, 2003; Chevance & Hughes, 2008; Macnab, 2003). The basal body contains a rotary motor that is powered by a transmembrane ion gradient and generates the torque needed for motility (Manson *et al*, 1977).

The torque-generating stator of the flagellar motor consists of a MotA pentamer and MotB dimer (Deme *et al*, 2020; Santiveri *et al*, 2020). MotB contains an N-terminal transmembrane domain, a plug/linker region, and a C-terminal periplasmic region that fixes the stator to the peptidoglycan layer through interactions with the peptidoglycan layer and the P ring protein FlgI (Hizukuri *et al*, 2010; Hizukuri *et al*, 2009). MotA has four transmembrane domains and together with MotB forms a proton channel in the inner membrane (Deme *et al.*, 2020; Santiveri *et al.*, 2020). In *Escherichia coli* and some other bacterial species, stator units are not permanently fixed at the motor but instead engage and disengage with the motor from a pool of unbound stators in the membrane (Leake *et al*, 2006; Tipping *et al*, 2013). In *E. coli*, the number of stators engaged within the motor varies according to the load, ranging from one or two under very low loads to an estimated 6-11 stators at high loads (Lele *et al*, 2013). In contrast to the *E. coli* motor, stator units are stably associated with the motors of *H. pylori* and some other bacterial species (Beeby *et al*, 2016; Guo *et al*, 2022; Qin *et al*, 2017; Tachiyama *et al*, 2022). When not engaged with the motor, the MotB plug blocks proton flow through the stator (Hosking *et al*, 2006). Following incorporation of the stator into the motor, MotB undergoes a conformation change that unplugs the stator and promotes dimerization of the periplasmic domain to allow binding of the protein to peptidoglycan (Hu *et al*, 2022). Proton flow through the stator is thought to power rotation of the MotA pentamer around the stationary MotB dimer (Deme *et al.*, 2020; Santiveri *et al.*, 2020). Engagement of MotA with the C ring protein FliG results in the rotation of the C ring/MS ring rotor complex, and the resulting torque is transmitted through the rod and hook to the filament, which acts as a propeller to push the cell forward (Blair & Berg, 1990; Stolz & Berg, 1991).

FliL is an integral membrane protein that is required for flagellar motor function in various bacteria (Attmannspacher *et al*, 2008; Jenal *et al*, 1994; Mengucci *et al*, 2020; Motaleb *et al*, 2011; Suaste-Olmos *et al*, 2010; Tachiyama *et al.*, 2022; Takekawa *et al*, 2019; Zhu *et al*, 2015). Cryo-electron tomography (cryo-ET) reconstructions of the *Borrelia burgdorferi* flagellar motor suggested FliL is part of the motor (Motaleb *et al.*, 2011). Recent cryo-ET studies of flagellar motors in *H. pylori* and *B. burgdorferi* showed the C-terminal periplasmic domain of FliL (FliL_C_) forms a ring that surrounds the plug/linker regions of the MotB dimer in its extended, active form (Guo *et al.*, 2022; Tachiyama *et al.*, 2022). Crystallographic studies of FliL_C_ proteins from *Vibrio alginolyticus* and *H. pylori* revealed a fold similar to that of the stomatin/prohibitin/flotillin/HflK/C (SPFH) domain of mouse stomatin (Tachiyama *et al.*, 2022; Takekawa *et al.*, 2019). SPFH family proteins often have roles in modulating the activities of ion channels (Cullinan *et al*, 2021), and FliL may have a similar function in modulating stator activity.

*Campylobacter jejuni*, a member of the phylum Campylobacterota, is closely related to *H. pylori*. In a high-throughput genetic screen of *C. jejuni* transposon mutants, Gao and co-workers identified several previously uncharacterized genes required for invasion of cultured mammalian cells and wild-type motility (Gao *et al*, 2014). The product of one of these genes, designated FlgV, interacted with the MS ring protein FliF in a co-immunoprecipitation (co-IP) assay (Gao *et al.*, 2014). We report here that *H. pylori* and other members of the phylum Campylobacterota have FlgV homologs. To examine the role of *H. pylori flgV* in flagellar function, we deleted *flgV* in *H. pylori* B128 and a highly motile variant of *H. pylori* G27 that we isolated and designated as strain G27M. In both B128 and G27M, deleting *flgV* resulted in a reduction in the number of flagella per cell and impaired motility of the strains in soft agar medium. Cryo-ET of *H. pylori* wild-type and Δ*flgV* mutants indicated FlgV forms a ring located inside the C ring and near the junction of the MS and the C rings. The location of the FlgV ring and reduced motility of the Δ*flgV* mutants in soft agar medium suggests roles for the FlgV ring in promoting rotor-stator interactions or switching of rotor direction. We hypothesize the FlgV ring is an adaptation that evolved to allow the large-diameter, high torque-generating motors of members of the phylum Campylobacterota to operate efficiently.

## RESULTS

### FlgV homologs are widespread among members of the phylum Campylobacterota

*H. pylori* 26695 was reported to lack a FlgV homolog (Gao *et al.*, 2014), but this report was based on an incorrect annotation of the *flgV* locus that identified an open reading frame in the opposite orientation (Tomb *et al*, 1997). A subsequent analysis of the region revealed an open reading frame between *flhG* and *fliA* that was in the same orientation as these genes, and the gene was given the locus tag HP1033b (Figure 1A) (Niehus *et al*, 2004). The revised annotation of the *flhG-fliA* region was consistent with the genomes of other *H. pylori* strains, including *H. pylori* G27, where HPG27_395 is the HP1033b homolog. A blastp analysis revealed homology between HPG27_395 and *C. jejuni* FlgV, although the sequence similarity between the two proteins is low (Table 1). HPG27_395 and *C. jejuni* FlgV are of similar lengths (104 and 118 amino acid residues in length, respectively), and both are encoded in genes located between *flhG* and *fliA* (Figure 1B). FlhG is a MinD-like protein that controls flagellum number in many polar-flagellated bacteria (Kojima *et al*, 2020; Schuhmacher *et al*, 2015), while FliA (σ^28^) is an alternative sigma factor required for transcription of flagellar genes whose products are required late in the flagellum assembly pathway (Aldridge & Hughes, 2002; McCarter, 2006; Smith & Hoover, 2009). HPG27_395 and *C. jejuni* FlgV are predicted to be integral membrane proteins with similar membrane topologies (Figure 1C). In addition, the tertiary structures of HPG27_395 and *C. jejuni* FlgV predicted by the AlphaFold tool in ChimeraX (Jumper *et al*, 2021) are very similar (Figure 1D). Taken together, these data provide strong evidence that HPG27_395 and *C. jejuni* FlgV are orthologs, and hence we refer to HPG27_395 as FlgV. A search of the genomes of representative genera of the phylum Campylobacterota revealed homologs of *C. jejuni* FlgV are widespread among members of the phylum (Table 1), and in each case, the *flgV* homolog is located between *flhG* and *fliA* (Figure 1B). Species of Campylobacterota in which we were unable to identify FlgV homologs by blastp included *Arcobacter butzleri*, *Thioreductor miscantisoli*, *Nitratifractor salsuginis,* and *Nitratiruptor tergarcus*. In the *A. butzleri* genome, however, there is a gene immediately downstream of *flhG* that encodes a protein of similar size, predicted membrane topology, and predicted tertiary structure to that of *C. jejuni* FlgV (Figure 1D), and we propose the *A. butzleri* protein is a FlgV homolog.

**Figure 1.**
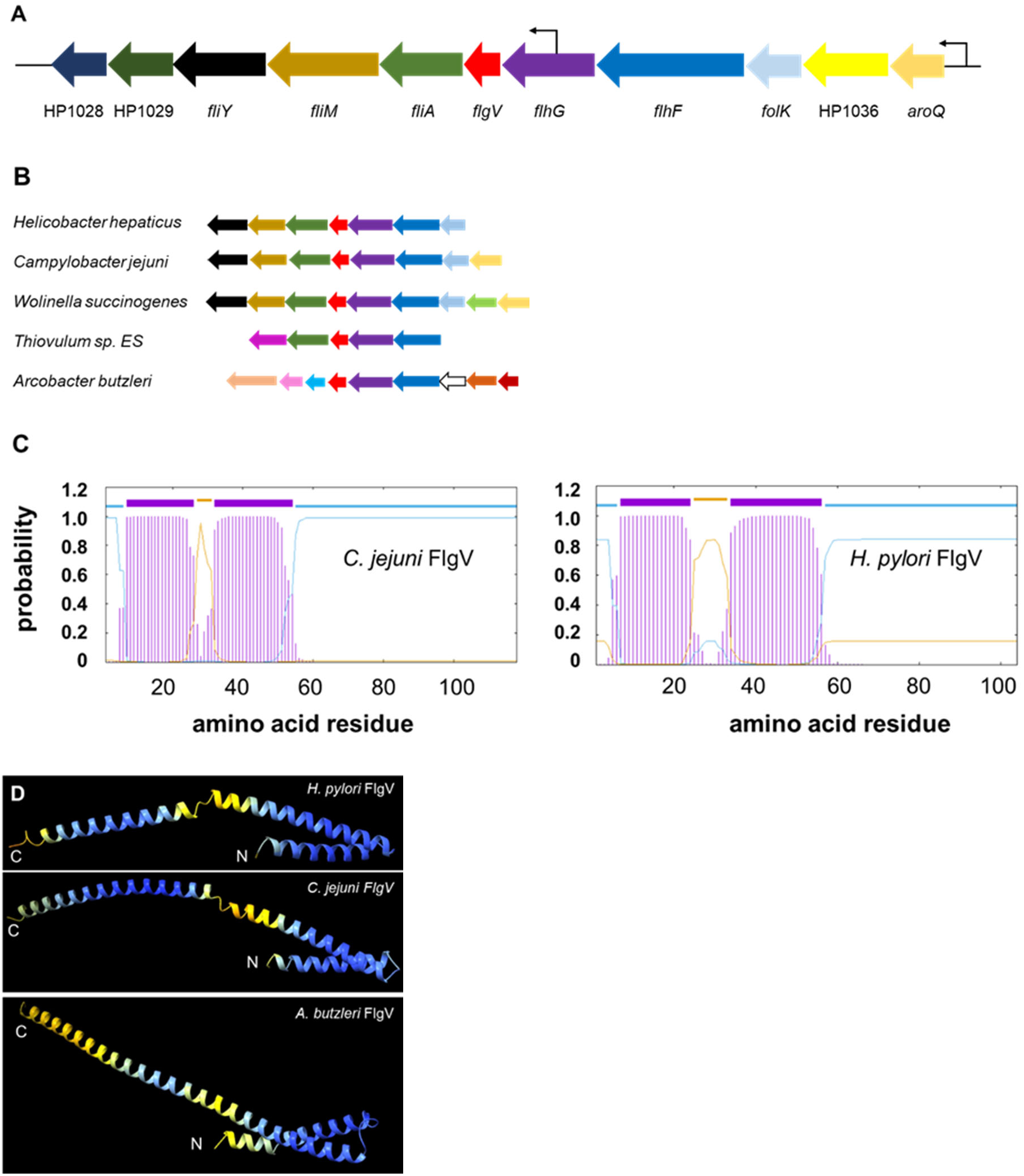
Organization of operons containing *flgV* in *H. pylori* and representative members of the phylum Campylobacterota. (**A**) Organization of genes within operon containing *flgV* in *H. pylori* 26695. The region corresponding to *flgV* (red) was annotated originally as an open reading frame (ORF) in the opposite direction, but reassessment of the region between *flhF* and HP1028 identified a new ORF (HP1033b) in the same orientation as the flanking ORFs (Niehus *et al.*, 2004). The smaller arrows upstream of *aroQ* and within *flhG* indicate identified transcriptional start sites in *H. pylori* 26695 (Sharma *et al*, 2010). Known flagellar genes in the operon include *flhF* (dark blue), *flhG* (purple), *fliA* (green), *fliM* (dark gold), and *fliY* (black). Other genes in the operon include *aroQ* (gold), *folK* (light blue), and three genes of unknown function (HP1036, yellow; HP1029, dark olive; and HP1028, navy blue). (**B**) Synteny of *flhFGflgV* in representative genera of Campylobacterota. Homologous genes are color coded as indicated in panel A. Genes that are not present in the *H. pylori* 26695 *flhFGflgV* operon include *pepQ* (light green), potential *flgJ* homolog (dark red), potential *flgN* homolog (dark orange), hypothetical protein (white with black outline), *fliE* (sky blue), *flgB* (pink), and *fliK* (peach). The gene organization around the *flhFGflgV* region in *Sulfurospirillum multivorans*, *Hydrogenimonas thermophila*, *Caminibacter mediatlanticus*, and *Nautilia profundicola* is similar to that of *W. succinogenes*. The gene organization around the *flhFGflgV* region in *Sulfurimonas autotrophica*, *Sulfuricurvum kujiense*, and *Lebetimonas natshushimae* is similar to that of *C. jejuni*. In *A. butzleri*, *flhFGflgV* appear to be part of a larger operon of ~30 genes encoding many known flagellar and chemotaxis proteins. (**C**) Predicted membrane topology of *C. jejuni* and *H. pylori* FlgV homologs. The amino acid sequences of *C. jejuni* 81-176 and *H. pylori* G27 FlgV homologs were analyzed using TMHMM - 2.0 (https://services.healthtech.dtu.dk/service.php?TMHMM-2.0), an online program for predicting transmembrane helices in proteins. On the top of the figures, predicted transmembrane helices are indicated by purple bars, regions of the protein predicted to be exposed on the cytoplasmic side of the membrane are indicated by blue lines, and regions of the protein predicted to be exposed on the periplasmic side of the membrane are indicated in gold lines. (**D**) Tertiary structures of FlgV proteins from *H. pylori*, *C. jejuni*, and *A. butzleri* predicted by the AlphaFold tool in ChimeraX (Jumper *et al.*, 2021) suggested the proteins consist primarily of α-helices. N-termini (N) and C-termini (C) of FlgV proteins are indicated. Regions in blue indicate a high confidence for predicted tertiary structure, while regions in yellow indicate a lower confidence for predicted tertiary structure.

**Table 1.** *C. jejuni* FlgV homologs in various genera of Campylobacterota identified by BlastP analysis.

| Description | Query coverage | Percent identity | Length | E value | Accession |
| --- | --- | --- | --- | --- | --- |
| hypothetical protein [*Campylobacter jejuni* 81-176] | 100% | 100.00% | 118 | 5e-80 | WP_002868772.1 |
| hypothetical protein [*Helicobacter pylori* G27] | 78% | 30.11% | 104 | 1e-07 | WP_000868000.1 |
| hypothetical protein [*Helicobacter hepaticus* ATCC 51449] | 43% | 43.14% | 118 | 1e-09 | WP_011115985.1 |
| Hypothetical protein [*Wolinella succinogenes* DSM 1740] | 94% | 27.59% | 117 | 2e-09 | WP_011139455.1 |
| hypothetical protein [*Sulfurimonas autotrophica* DSM 16294] | 86% | 38.24% | 111 | 1e-05 | WP_013326520.1 |
| hypothetical protein [*Sulfurospirillum multivorans* DSM 12446] | 91% | 41.05% | 124 | 5e-15 | WP_025343576.1 |
| hypothetical protein [*Sulfuricurvum kujiense* DSM 16994] | 75% | 30.34% | 107 | 3e-11 | WP_013459341.1 |
| hypothetical protein [*Hydrogenimonas thermophila*] | 75% | 33.71% | 109 | 1e-09 | WP_092910690.1 |
| hypothetical protein ThvES_00009150 [*Thiovulum* sp. ES] | 84% | 28.95% | 112 | 4e-04 | EJF07001.1 |
| hypothetical protein [*Caminibacter mediatlanticus* TB-2] | 77% | 40.82% | 112 | 7e-08 | WP_007473940.1 |
| hypothetical protein [*Lebetimomas natshushimae*] | 77% | 39.13% | 112 | 2e-11 | WP_096259429.1 |
| hypothetical protein [*Nautilia profundicola* Am-H] | 77% | 36.17% | 111 | 2e-09 | WP_012663466.1 |
| hypothetical protein [*Arcobacter butzleri* 7h1h] | <sup>a</sup>nssf | <sup>a</sup>nssf | 127 | >5e-03 | WP_014469277.1 |

<sup>a</sup>No significant similarity found

### FlgV is required for robust motility of *H. pylori* in soft agar medium

To determine if FlgV was required for flagellum function in *H. pylori*, we initially deleted *flgV* in *H. pylori* G27 and examined the motility of the resulting mutant in soft agar medium. Although the Δ*flgV* mutant produced a smaller swim halo than the G27 parental strain, the parental strain produced a relatively small swim halo, which limited our confidence in ascribing a motility defect to the Δ*flgV* mutant. To address this issue, we isolated a highly motile variant of *H. pylori* G27 from a motility enrichment and designated the motile variant as G27M. Whole genome sequencing of G27M revealed it had 80 single-nucleotide polymorphisms (SNPs) and insertion/deletions (indels) compared to the *H. pylori* G27 genome sequence in the GenBank nucleotide sequence database (Tables S1 and S2). Most of these SNPs and indels had presumably accumulated during repeated passage of the strain in the laboratory and had no role in the enhanced motility of G27M. The vast majority of the SNPs and indels were in pseudogenes or intergenic regions (51 and 20 SNPs/indels, respectively; Tables S1 and S2). Three of the SNPs though were in flagellar genes. Two of the flagellar gene SNPs were in *flgH*, which encodes the L ring protein, and changed Gly-178 to Cys (GGG to TGT; we designated the *flgH* allele as *flgH178*). The other flagellar gene SNP was a nonsense mutation in codon 78 of *fliL* (CAG to TAG; Gln-78 to stop; we designated allele as *fliL78*), which resulted in a predicted truncation of FliL at the boundary of the linker region and FliL_C_.

Deleting *flgV* in *H. pylori* G27M resulted in significant reductions in the motility of the strain in soft agar medium and the number of flagella per cell (Figure 2). Introducing *flgV* on the shuttle vector pHel3 into the Δ*flgV* mutant rescued motility and the number of flagella per cell (Figure 2), indicating loss of FlgV was responsible for the decreased motility and reduction in the number of flagella per cell in the Δ*flgV* mutant.

**Figure 2.**
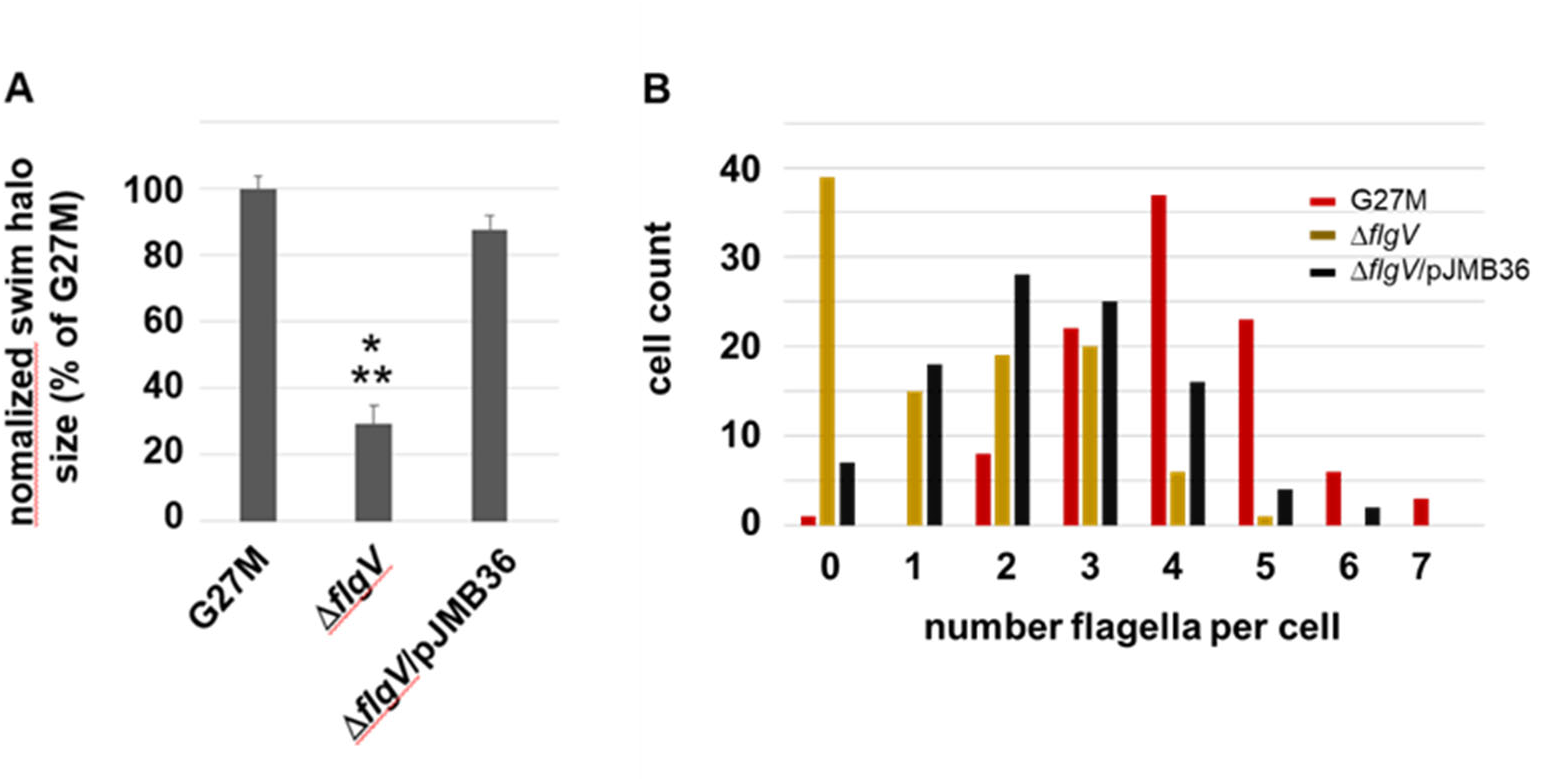
Motility and flagellation phenotypes of *H. pylori* G27M Δ*flgV* mutant. (**A**) Motility of *H. pylori* G27M, Δ*flgV* mutant, and Δ*flgV* complemented strains (Δ*flgV*/pJMB36). Strains were stab inoculated into soft agar medium, and diameters of the resulting swim halos were measured following 7 d incubation. Bars indicate mean values for swim halo diameters normalized to the mean swim halo value for *H. pylori* G27M. Number of technical replicates for the strains ranged from 3 to 12. Error bars indicate one standard deviation. Singe asterisk indicates swim halo diameters that differed significantly from G27M (*p*-value <0.00001). Double asterisk indicates the swim halo diameters of Δ*flgV* mutant differed significantly from that of the Δ*flgV* mutant that carried plasmid pJMB36 (*p*-value <0.00001). Statistical analysis of the data was done using a two-sample *t* test. (**B**) Flagella were counted for at least 100 cells for *H. pylori* G27M (red), G27M Δ*flgV* mutant (gold), and G27M Δ*flgV* mutant that carried the complementation plasmid pJMB36 (black). The complemented strain had significantly more flagella per cell than the original Δ*flgV* mutant (*p*-value <0.00001). Statistical significance for differences in the distribution of the number of flagella per cell were determined using a Mann-Whitney U test.

### Isolation of motile variants of the *H. pylori* G27M Δ*flgV* mutant

To identify mutations that by-pass the requirement of FlgV for robust motility, we enriched for variants of the Δ*flgV* mutant that had enhanced motility in soft agar medium. Three independent motile variants were isolated (designated MV1 through MV3) that had motilities in soft agar medium that approximated that of G27M (Figure 3A). Interestingly, all three of the motile variants possessed flagella that were substantially longer than the flagella of G27M (Figure 3B and 3C). Flagella of MV1 and MV3 were on average ~2.5 longer than the flagella of G27M (Table 2). About 25% of the MV2 flagella were longer than the longest flagella measured for G27M (>6 mm in length), but a large proportion of MV2 flagella were 2 μm or less in length (Figure 3C). Consequently, the mean flagellar length in MV2 did not differ significantly from that of G27M. Two of the motile variants, MV1 and MV2, produced significantly more flagella per cell than the Δ*flgV* parental strain, although flagella number was not restored to the level observed for G27M in these motile variants (Figure 3C). Cells of MV3 appeared to be slightly more flagellated than those of the Δ*flgV* parental strain, but the difference was just above the cutoff for statistical significance (*p*-value = 0.057).

**Figure 3.**
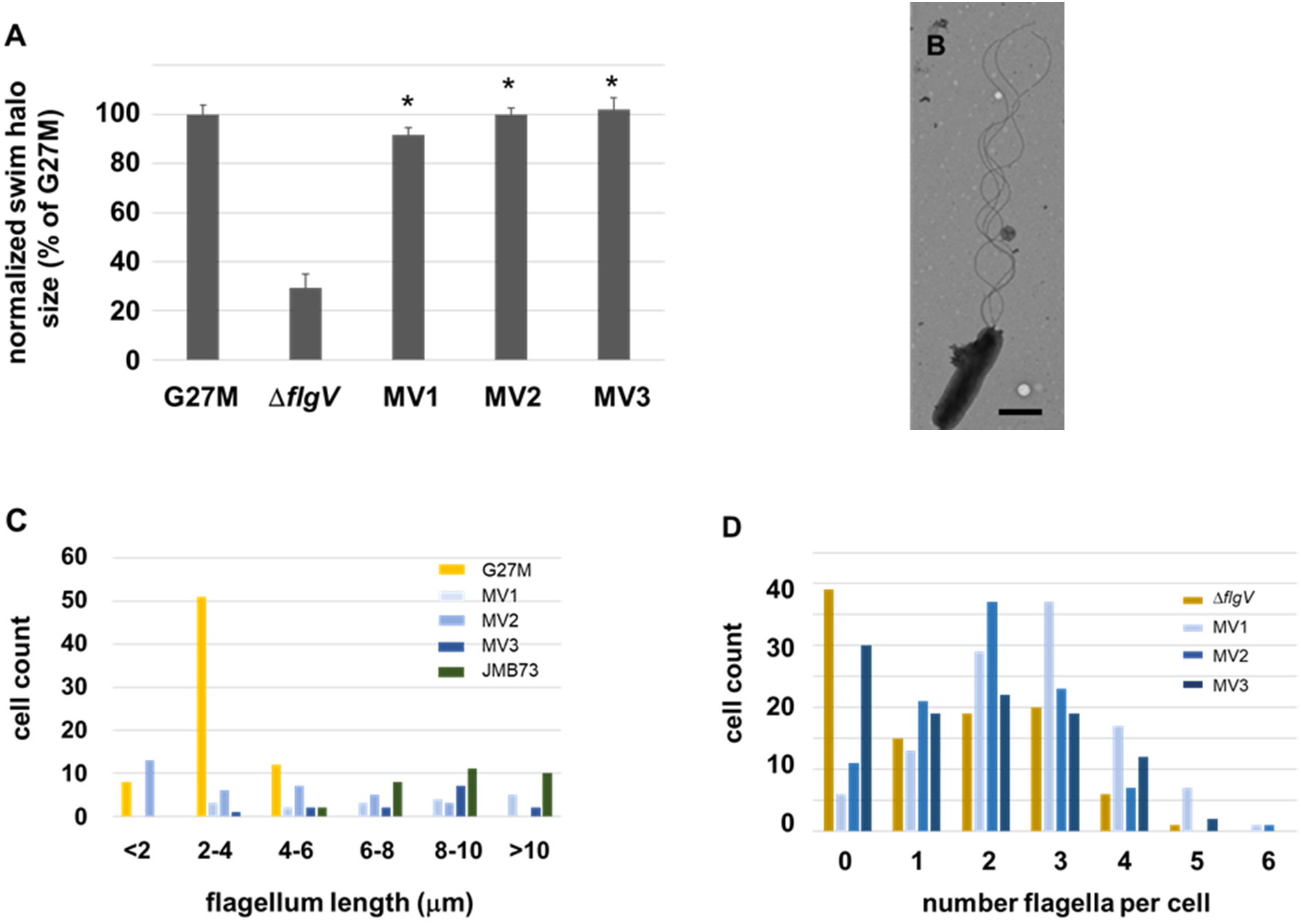
Motility and flagellation patterns of motile variants of the G27M Δ*flgV* mutant. (**A**) Motilities of the Δ*flgV* motile variants were compared with those of *H. pylori* G27M and the G27M Δ*flgV* parental strain in soft agar medium. Strains were stab inoculated into soft agar medium, and diameters of the resulting swim halos were measured following 7 d incubation. Bars indicate mean values for swim halo diameters normalized to the mean swim halo value for *H. pylori* G27M. The number of technical replicates for the strains ranged from 3 to 12. Error bars indicate one standard deviation. Asterisk indicates swim halo diameter that was significantly larger than that of the Δ*flgV* parental strain (*p*-value <0.00001). Statistical analysis of the data was done using a two-sample *t* test. (**B**) TEM image of a typical Δ*flgV* motile variant MV1 cell. Bar equals 1 μm. (**C**) Flagellum length distributions for the various strains. Flagella were grouped according to their measured lengths as indicated in x-axis. (**D**) Flagella were counted for at least 100 cells for *H. pylori* G27M Δ*flgV* parental strain (gold), and the Δ*flgV* motive variants - MV1 (light blue), MV2 (medium blue), and MV3 (dark blue). Strains MV1 and MV2 had significantly more flagella per cell the Δ*flgV* parental strain (*p*-values <0.00001 and 0.00066, respectively), whereas the distribution of the number of flagella per cell for strain MV3 and the Δ*flgV* parental strain did not differ significantly (p-value = 0.057). Statistical significance for differences in the distribution of the number of flagella per cell were determined using a Mann-Whitney U test.

**Table 2.** Lengths of flagella in *H. pylori* G27M and G27M Δ*flgV* motile variants.

| Strain | Median flagellum length ($\mu\text{m}$) | Mean flagellum length ($\mu\text{m}$)<sup>a</sup> | Number measured | *p*-value<sup>b</sup> |
| --- | --- | --- | --- | --- |
| G27M | 3.28 | 3.16 ± 0.96 | 71 | |
| MV1 | 8.72 | 7.61 ± 2.98 | 15 | <1x10<sup>-5</sup> |
| MV2 | 2.96 | 4.00 ± 2.70 | 34 | 0.78 |
| MV3 | 8.74 | 8.16 ± 2.30 | 14 | <1x10<sup>-5</sup> |
| JB73 | 9.47 | 9.49 ± 2.26 | 33 | <1x10<sup>-5</sup> |

<sup>a</sup> Range indicates one standard deviation.
<sup>b</sup> Significant differences in flagellum length for the motile variants compared to G27M were determined using a Mann-Whitney U test.

Whole-genome sequencing of the Δ*flgV* motile variants revealed all three strains contained a single-nucleotide polymorphism (SNP) or deletion in *flaG*-2 (HPG27_708) (Table 3). MV1 and MV3 had the same mutation in *flaG*-2, a C to T transition at nucleotide position 313 that changed the CAA codon (Gln-105) to a UAA stop codon, resulting in a 15 amino acid truncation of the protein. The *flaG*-2 mutation in MV2 was a deletion of a T at nucleotide position 102 that causes a frameshift resulting in a truncated protein with only the first 33 (out of 119) amino acid residues of FlaG-2. The mutations in *flaG*-2 were likely responsible for the longer flagella of the Δ*flgV* motile variants, as mutations in *flaG*-2 homologs in *Pseudomonas fluorescens* and *Vibrio anguillarum* result in abnormally long flagella (Capdevila *et al*, 2004; McGee *et al*, 1996)In addition to the lesion in *flaG*-2, each Δ*flgV* motile variant contained mutations in *faaA* (HPG27_570) and HPG27_125. FaaA is an autotransporter (Type V protein secretion pathway protein) that shares homology with the vacuolating toxin VacA and localizes to the flagellar sheath (Radin *et al*, 2013). MV1 and MV3 shared the same deletion in *faaA* (G residue at nucleotide position 623), while MV2 had a deletion of a G in a homopolymeric run of eight G residues at nucleotide position 4,493. All three motile variants had the same frameshift mutation in HPG27_125, which encodes a predicted iron-sulfur cluster-binding protein.

**Table 3.** Identified intragenic mutations in *H. pylori* G27M Δ*flgV* motile variants.

| Isolate | Gene description | Locus tag | <sup>a</sup>Sequence | <sup>b</sup>Impact | <sup>c</sup>Freq |
| --- | --- | --- | --- | --- | --- |
| MV1 | LutB/LldF family<br>L-lactate oxidation<br>iron-sulfur protein | HPG27_RS00700<br>(HPG27_125) | (C)6→7<br>coding (1199/1446 nt) | V399fs | 97.2% |
|  | *fliN* | HPG27_RS02835<br>(HPG27_543) | ACT→GCT | T28A | 98.7% |
|  | *faaA* | HPG27_RS02965<br>(HPG27_570) | Δ1 bp (627/9579 nt) | N209fs | 98.9% |
|  | *flaG-2* | HPG27_RS03655<br>(HPG27_708) | CAA→TAA | Q105* | 99.7% |
|  | *fliL* | HPG27_RS03950<br>(HPG27_765) | Δ1 bp (187/552 nt) | P63fs | 98.6% |
| MV2 | LutB/LldF family<br>L-lactate oxidation<br>iron-sulfur protein | HPG27_RS00700<br>(HPG27_125) | (C)6→7<br>coding (1199/1446 nt) | V399fs | 97.0% |
|  | *faaA* | HPG27_RS02965<br>(HPG27_570) | (G)8→7 (4493/9579 nt) | G1497fs | 96.9% |
|  | *flaG-2* | HPG27_RS03655<br>(HPG27_708) | Δ1 bp (102/360 nt) | I34fs | 98.0% |
|  | Jag N-terminal<br>domain-containing<br>protein | HPG27_RS07190<br>(HPG27_1372) | GCC→GTC | A50V | 97.4% |
| MV3 | LutB/LldF family<br>L-lactate oxidation<br>iron-sulfur protein | HPG27_RS00700<br>(HPG27_125) | (C)6→7<br>coding (1199/1446 nt) | V399fs | 96.9% |
|  | TrkA C-terminal<br>domain-containing<br>protein | HPG27_RS01415<br>(HPG27_261) | (A)9→8 (946/1437 nt) | M316fs | 97.2% |
|  | *faaA* | HPG27_RS02965<br>(HPG27_570) | Δ1 bp (627/9579 nt) | N209fs | 98.9% |
|  | *flaG-2* | HPG27_RS03655<br>(HPG27_708) | CAA→TAA | Q105* | 98.4% |
|  | *fliL* | HPG27_RS03950<br>(HPG27_765) | Δ1 bp (187/552 nt) | P63fs | 99.6% |

<sup>a</sup>The numbers in parentheses indicate the position of the mutation (first number) within the entire length of the open reading frame (second number).
<sup>b</sup>Indicates the position where a different amino acid was introduced, position where a stop codon was introduced (\*), or site where a frameshift mutation occurred (fs).
<sup>c</sup>Indicates the percentage of reads at the position that had the particular SNP.

Each Δ*flgV* motile variant possessed one or two additional mutations in coding sequences, and two of these mutations were in known flagellar genes. MV1 and MV3 had the same deletion in *fliL*, which introduced a frameshift mutation in codon 63 (Table 3). MV1 also had a SNP in *fliN* (HPG27_543; encodes a C ring protein) that resulted in an alanine substitution at Thr-29. In addition to the mutations within coding regions of genes, the *flgV* motile variants had two to five mutations in intergenic regions (Table S3), though none of the intergenic regions with mutations were adjacent to known flagellar genes.

Since all three of the motile variants had mutations in *flaG*-2, *faaA* and HPG27_125, we reasoned one or more of these mutations were responsible for rescuing motility in the Δ*flgV* mutant. HPG27_125 encodes a predicted FeS protein that likely has a role in metabolism, and we considered it unlikely the mutation in this gene was responsible for rescuing motility in the Δ*flgV* mutant. Given that FaaA localizes to the flagellar sheath (Radin *et al.*, 2013) and FlaG-2 controls flagellum length (Capdevila *et al.*, 2004; McGee *et al.*, 1996), the mutations in either *faaA* or *flaG*-2 seemed the most likely candidates for rescuing motility in the Δ*flgV* mutant. Introduction of either the *faaA* allele from MV2 (designated *faaA4493*) or the *flaG*-2 allele from MV1 (designated *flaG-2313*) into the G27M Δ*flgV* parental strain, however, failed to rescue motility in the strain. Introducing the *flaG-2313* allele into the G27M *flgV* mutant (strain JB73), however, did result in longer flagella (Figure 3C; Table 2), which confirmed the allele was responsible for increased flagellar length in MV1 and MV3. It is possible that mutations in both *flaG*-2 and *faaA* were needed to rescue motility in the G27M Δ*flgV* mutant. We did not pursue this line further by introducing the *faaA4493* and *flaG-2313* alleles together into the G27M Δ*flgV* mutant, as motile variants of a *H. pylori* B128 Δ*flgV* mutant did not have mutations in either *faaA* or *flaG*-2 (Table S4).

### Analysis of conserved amino acid residues in *H. pylori* FlgV

An alignment of the amino acid sequences of FlgV homologs from a diverse set of bacterial species within the phylum Campylobacterota revealed a few conserved amino acid positions (Figure S1). Most notable of the conserved positions is a GFF motif within the first predicted transmembrane helix of FlgV. To determine the importance of conserved amino acid residues for FlgV function, we changed two pairs of adjacent amino acid residues (Phe-15/Phe-16 and Glu-71/Glu-72) to alanine residues and examined the ability of the resulting FlgV variants to rescue motility in the G27M Δ*flgV* mutant. Both the FlgV^F15A,F16A^ variant and the FlgV^E71A,E72A^ variant supported robust motility of the G27M Δ*flgV* mutant in soft agar medium (Figure 4A). The strain expressing the FlgV^E71A,E72A^ variant was highly flagellated and had significantly more flagella per cell than the G27M Δ*flgV* parental strain (Figure 4B). In contrast, the strain expressing the FlgV^F15A,F16A^ variant was poorly flagellated and about half the cells examined were aflagellated (Figure 4B). Thus, even though a high proportion of the cells of the strain expressing the FlgV^F15A,F16A^ variant lacked flagella, enough of the cells were motile to generate a large swim halo. Taken together, these data suggest amino acid residues Phe-15 and Phe-16 in FlgV are important for flagellum assembly or stability but are not critical for flagellum function.

**Figure 4.**
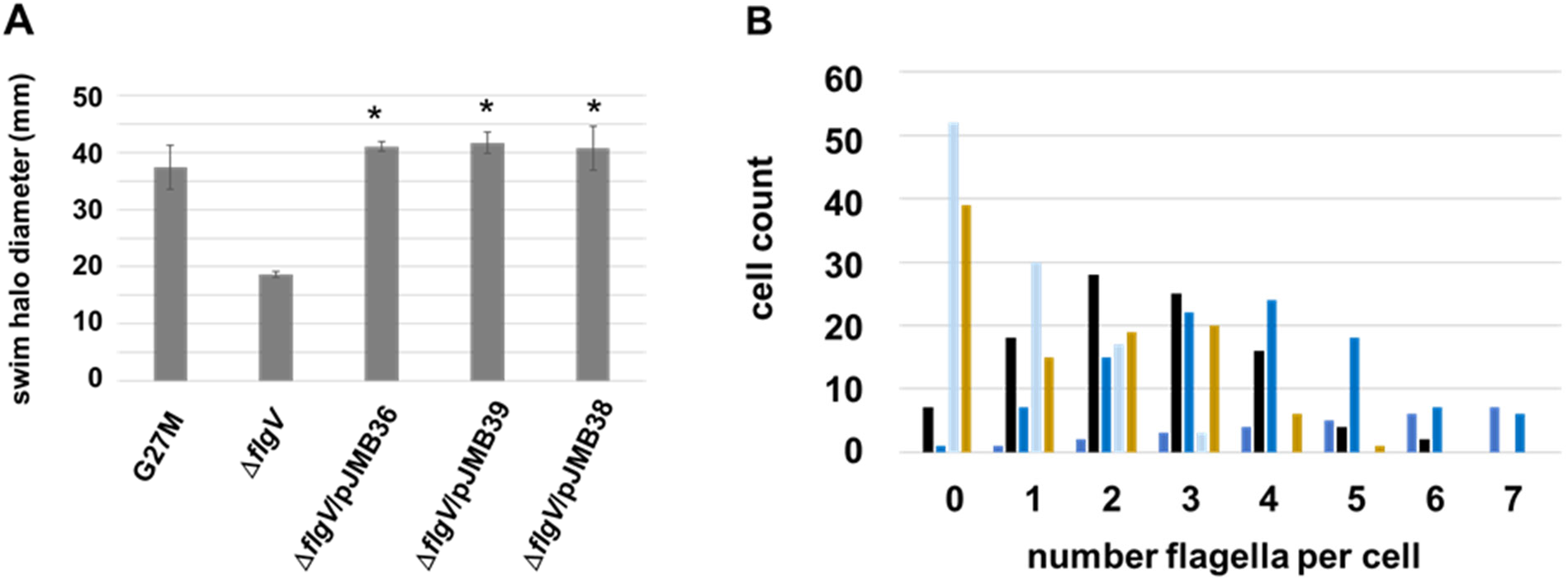
Motility and flagellation phenotypes of *H. pylori* G27M expressing FlgV variants. (**A**) Motilities of *H. pylori* G27M Δ*flgV* expressing FlgV^F15A,F16A^ variant (Δ*flgV*/pJMB39) or FlgV^E71A,E72A^ variant (Δ*flgV*/pJMB38) were compared with the motilities of *H. pylori* G27M (G27M), *H. pylori* G27M Δ*flgV* parental strain (Δ*flgV*), and G27M Δ*flgV* strain complemented with wild-type *flgV* (Δ*flgV*/pJMB36) in soft agar medium. Strains were stab inoculated into soft agar medium, and diameters of the resulting swim halos were measured following 7 d incubation. Bars indicate mean values for swim halo diameters for three technical replicates for each strain. Error bars indicate one standard deviation. Asterisk indicates swim halo diameter that was significantly larger than that of the Δ*flgV* parental strain (*p*-value <0.00001). Statistical analysis of the data was done using a two-sample *t* test. (**B**) Flagella were counted for at least 100 cells for *H. pylori* G27M Δ*flgV* parental strain (gold) and *H. pylori* G27M Δ*flgV* expressing FlgV (black), FlgV^E71A,E72A^ variant (dark blue), or FlgV^F15A,F16A^ variant (light blue). The strain expressing the FlgV^E71A,E72A^ variant, but not the strain expressing the FlgV^F15A,F16A^ variant, had significantly more flagella per cell than the Δ*flgV* parental strain (*p*-value <0.00001). Statistical significance for differences in the distribution of the number of flagella per cell were determined using a Mann-Whitney U test.

### FlgV forms a flagellar motor ring closely associated with the MS and C rings

The motility defect of the *H. pylori* G27M Δ*flgV* mutant suggests the motor of the mutant does not function properly in soft agar medium. To determine if this reduced function is due to a structural flaw, we compared the *in-situ* flagellar motor structures of wild-type *H. pylori* G27 and the *H. pylori* G27M Δ*flgV* mutant, using cryo-ET and subtomogram averaging. The motor of wild-type *H. pylori* G27 had a globular density within the C ring near its junction with the MS ring (Figure 5A), and this density was absent from the *H. pylori* G27M Δ*flgV* motor (Figure 5C). The electron density in the wild-type *H. pylori* G27 motor corresponded to a novel ring structure located inside the C ring (Figure 5E) that was absent from the motor of the *H. pylori* G27M Δ*flgV* mutant (Figure 5G). Moreover, the motor of the *H. pylori* G27M Δ*flgV* mutant also lacked the FliL_C_ ring (Figure 5K), in agreement with results from the genome sequence of *H. pylori* G27M that predicted the strain does not express a full-length FliL protein bearing the FliL_C_ domain (Table S1).

**Figure 5.**
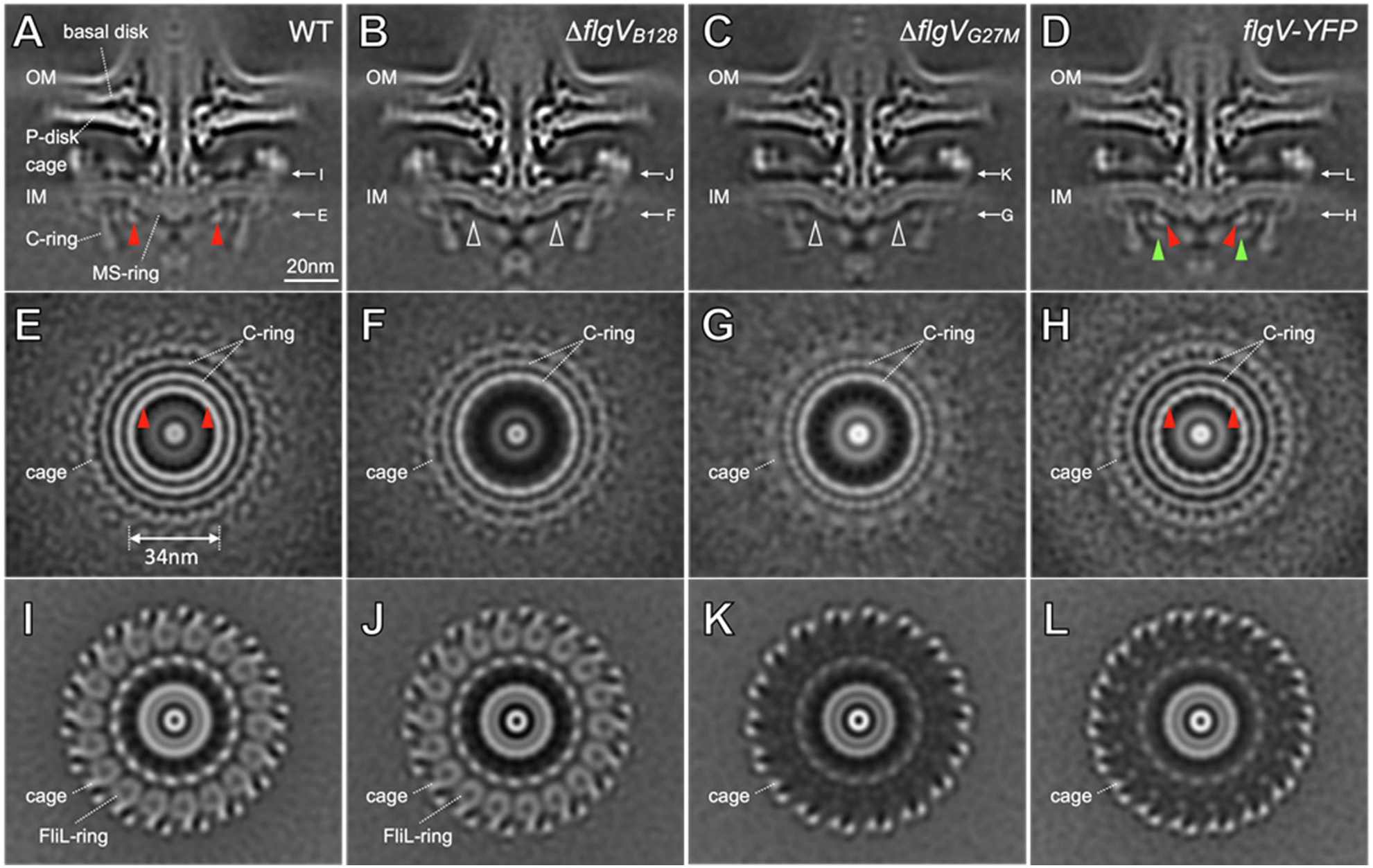
*In-situ* structures of the flagellar motor in *H. pylori* G27 wild-type and *flgV* mutant cells. (**A**-**D**) Medial slices through *in-situ* structures of G27 WT and several *flgV* mutant motors determined by cryo-ET and subtomogram averaging, respectively. A novel ring-like structure (highlighted by red arrows) in the G27 WT motor (**A**) is absent as shown by white arrows in the B128 Δ*flgV* mutant (Δ*flgV*_B128_) (**B**) and G27M Δ*flgV* mutant (Δ*flgV*_G27M_) (**C**). (**D**) Extra densities highlighted by green arrows are adjacent to the ring-like structure (red arrows) in G27M expressing FlgV-YFP. (**E**-**H**) Perpendicular cross-sections at the top of the C ring through the G27 WT and *flgV* mutant motor structures corresponding to the panels (**A-D**), respectively. Note the absence of electron densities corresponding to the FlgV ring in two Δ*flgV* mutants (**F, G**). (**I**-**L**) Different perpendicular cross-sections at the FliL ring level through the WT and *flgV* mutant motor structures corresponding to the panels (**A-D**), respectively. Note the absence of the FliL ring in the motors of the two mutants derived from strain G27M (**K, L**). OM, outer membrane; IM, inner membrane.

The absence of the novel motor ring in the Δ*flgV* mutant strongly suggests that this ring is formed by FlgV. To obtain direct evidence that FlgV does indeed form this previously unidentified motor ring, we expressed a FlgV-yellow fluorescent protein (YFP) fusion protein from the native *flgV* locus in *H. pylori* G27M. The motility of the strain expressing the FlgV-YFP fusion protein in soft agar medium did not differ significantly from that of G27M (Figure S2). Cryo-ET subtomogram reconstructions of intact motors of the *H. pylori* G27M strain expressing the FlgV-YFP fusion protein revealed that the identified ring is present (Figure 5H) and that an additional globular density is associated with the previously uncharacterized motor ring (Figure 5D). This additional density is likely due to the YFP that was linked to FlgV. Taken together, these data indicate FlgV forms a motor ring – designated here as the FlgV ring – that is closely associated with the MS and C rings.

Comparing the *in-situ* structures of the motors of the G27M Δ*flgV* mutant and G27M strain expressing the FlgV-YFP fusion showed the stator units were clearly visible in the latter strain but not in the G27M Δ*flgV* mutant (Figures S4G and S4H). Similarly, stator units were clearly visible in the motor of strain JMB79, which is the G27M Δ*flgV* mutant complemented with *flgV* (Figure S4E). The motor of the G27M Δ*flgV* mutant did contain the cage surrounding the motor (Figures 5C, 5G, and 5K), indicating that formation of the cage does not require the FlgV ring, FliL_C_ ring, or stator units.

### FlgV ring contributes to motility in *H. pylori* B128

As the G27M motor lacks the FliL_C_ ring, we wished to determine if the phenotype of the Δ*flgV* mutant is dependent on the absence of FliL_C_. Therefore, we deleted *flgV* from *H. pylori* B128, which expresses a full-length FliL protein, and characterized the resulting mutant. As observed with the G27M Δ*flgV* mutant, the B128 Δ*flgV* mutant generated smaller swim halos and significantly fewer flagella per cell compared to wild type (Figures S3A and S3B). The *in-situ* structure of the B128 Δ*flgV* mutant confirmed the absence of a FlgV ring (Figures 5B and 5F). As expected, the FliL_C_ ring was evident in the B128 Δ*flgV* motor (Figures 5B and 5J). In contrast to the G27M Δ*flgV* mutant motor, stator units were present in the motor of the B128 Δ*flgV* mutant (Figure S4B and S4F). Taken together with the data for the motors of the G27M strain expressing the FlgV-YFP fusion (Figure S4H) and the G27M Δ*flgV* complemented strain (Figure S4E), these results suggest that the combined loss of the FlgV ring and FliL_C_ ring prevents stable association of stator units with the motor.

As with the G27M Δ*flgV* mutant, we enriched for variants of the B128 Δ*flgV* mutant that had enhanced motilities in soft agar medium. Five clonal isolates of the B128 Δ*flgV* motile variants were obtained from three independent enrichments. Diameters of swim halos formed by clonal isolates of the B128 Δ*flgV* motile variants following 7-d incubation ranged from 24 to 32 mm, compared to ~10 mm for the parental B128 Δ*flgV* mutant. Whole-genome sequencing revealed that the B128 Δ*flgV* motile variants had two to four intragenic mutations (Table S4), none of which were in genes that had mutations in the G27M Δ*flgV* motile variants (Table 3). Each of the B128 Δ*flgV* motile variants had a deletion of a CT dinucleotide in *oipA* (encodes outer membrane inflammatory protein A; Table S4). The deleted CT dinucleotide is part of a hypermutable CT dinucleotide repeat motif within the 5’-region of *oipA* that is involved in regulating expression of the gene by slipped-strand mispairing (Miftahussurur & Yamaoka, 2015). The mutation in *oipA* was not likely responsible for rescuing motility in the B128 Δ*flgV* motile variants because the *H. pylori* B128 parental strain used to construct the Δ*flgV* mutant lacked one of the hypermutable CT dinucleotide repeat motifs in *oipA*, and the motile variants lost a second CT dinucleotide repeat. Thus, both the original B128 Δ*flgV* mutant and the B128 Δ*flgV* motile variants have frameshift mutations in *oipA*. Zero to three intergenic mutations were identified in the B128 Δ*flgV* motile variants, and none of the intergenic mutations were adjacent to known flagellar genes (Table S5). While we were unable to identify the genetic basis for the suppression of the motility defects in either the B128 or G27M Δ*flgV* mutants, future studies may reveal the mechanisms underlying how the *H. pylori* motor uses alternative pathways to rescue bacterial motility in the absence of the FlgV ring.

## DISCUSSION

The *H. pylori* flagellar motor contains the core components found in the archetypical motors of *E. coli* and *Salmonella enterica* (e.g., MS ring, C ring, L ring, P ring, rod, and MotA/MotB stator), as well as additional accessories not found in the *E. coli* and *S. enterica* motors (Qin *et al.*, 2017). As in the *H. pylori* motor, the flagellar motors of many other bacteria have accessories not found in the *E. coli* and *S. enterica* motors (Beeby *et al.*, 2016; Chaban *et al*, 2018; Chang *et al*, 2019; Zhu *et al*, 2019). We report here on a previously uncharacterized *H. pylori* motor accessory formed by FlgV. The FlgV ring is located in the interior of the C ring near the junction of the MS and C rings (Figures 5A, 5D, and 6). Consistent with the observed location of the FlgV ring, Gao and co-workers reported interactions between FliF and FlgV in a co-immunoprecipitation assay (Gao *et al.*, 2014). FlgV homologs are found in many members of the phylum Campylobacterota (Table 1), and we expect the motors of these bacteria to contain the FlgV ring.

**Figure 6.**
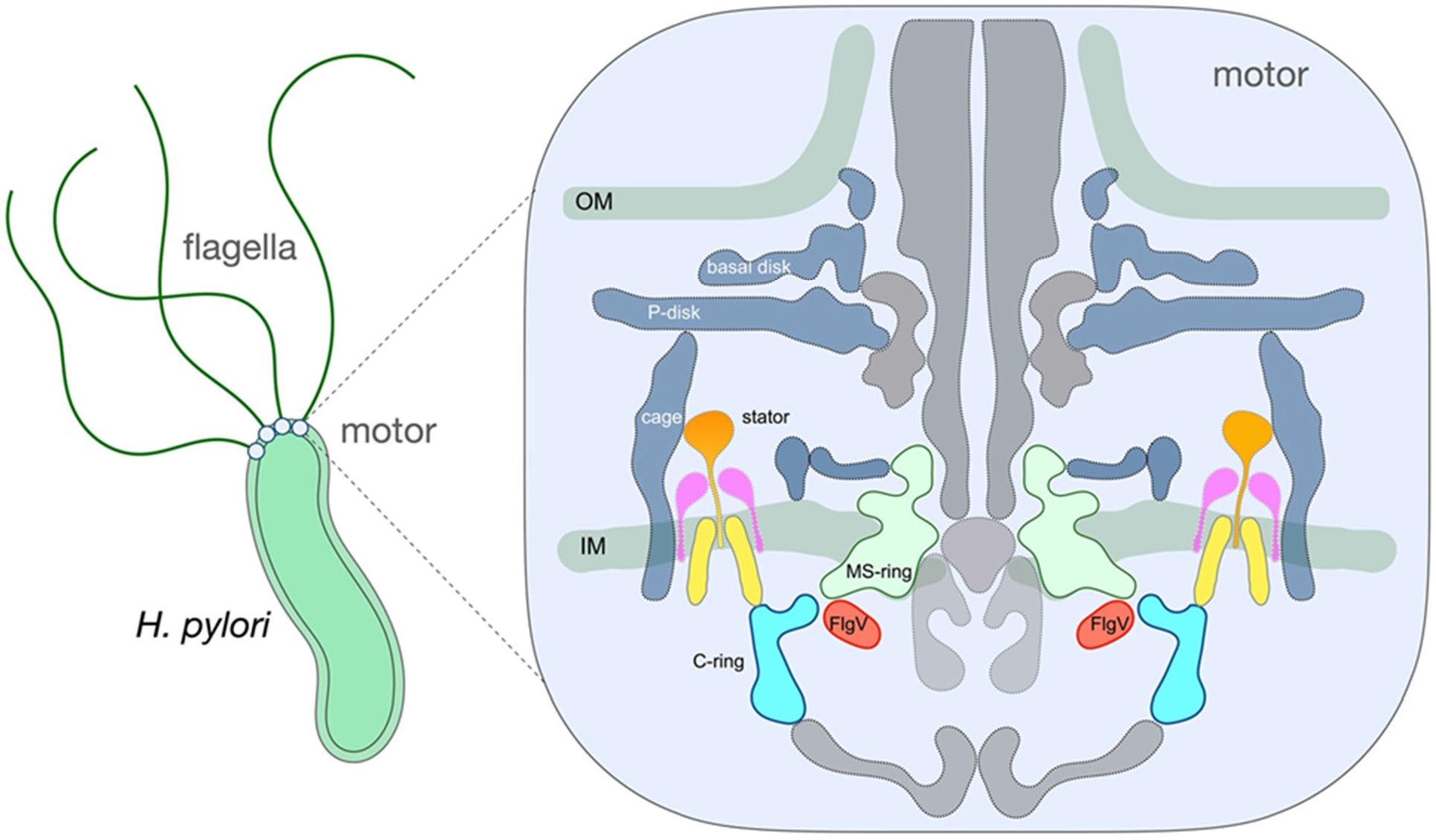
A model of the *H. pylori* motor. The FlgV ring (red) is located within the interior of the C ring (light blue) and interacts with the junction between the C ring and the MS ring (light green). FlgV closely associates with C ring protein FliG and MS ring protein FliF. Core motor components (flagellar protein export apparatus, rod, hook, P ring, and L ring) are shown in gray. Stator proteins MotB and MotA are shown in orange and yellow, respectively. FliL is shown in pink. *H. pylori* motor accessories are shown in dark blue and include the cage-like structure, P disk, and basal disk. Outer membrane (OM); inner membrane (IM).

The Δ*flgV* mutants in both the G27M and B128 backgrounds had fewer flagella compared to their parental strains (Figures 2B and S2B), indicating a potential role for FlgV in assembly or stability of the flagellum. In support of a possible role for FlgV in flagellum assembly or stability was the observation the FlgV^F15A,F16A^ variant supported robust motility in the G27M Δ*flgV* mutant, but failed to improve the flagellation of the strain (Figure 4). These findings suggest FlgV has distinct and separable roles in motor function and flagellum assembly or stability.

Given the synteny of *flhFG* and *flgV* in members of the phylum Campylobacterota (Figure 1B), FlgV may work with FlhF or FlhG to control the flagellation pattern in *H. pylori*. FlhF and FlhG control the flagellation pattern in many bacteria that have polar flagella (Schuhmacher *et al.*, 2015). FlhF is a GTPase that cycles between a GTP-bound form that facilitates flagellum assembly at the cell pole, and an inactive GDP-bound or apo-form (Schuhmacher *et al.*, 2015). Mutations in *flhF* typically result in a significant reduction in the proportion of flagellated cells and increased localization of flagella to nonpolar sites (Balaban *et al*, 2009; Green *et al*, 2009; Murray & Kazmierczak, 2006; Pandza *et al*, 2000). FlhG controls flagella number by modulating the GTPase activity of FlhF, and mutations in *flhG* generally result in hyperflagellation (Balaban & Hendrixson, 2011; Campos-Garcia *et al*, 2000; Correa *et al*, 2005; Dasgupta *et al*, 2000; Kusumoto *et al*, 2006). Although we did not observe the improper localization of flagella to nonpolar sites in the Δ*flgV* mutants, the reduced number of flagella in the Δ*flgV* mutants may indicate that FlgV works with FlhF to initiate flagellum assembly at the cell pole.

The MS ring and C ring form the rotor of the flagellar motor. The two rings are linked together by interactions between the C-terminal, cytoplasmic domain of FliF and the N-terminal domain of FliG (Lynch *et al*, 2017; Xue *et al*, 2018). The transfer of torque from the stator unit to the rotor involves electrostatic interactions between conserved acidic and basic residues in MotA and the FliG Helix_Torque_, an α-helix in the C-terminal domain of the protein (Santiveri *et al.*, 2020). Given the proximity of the FlgV ring to the junction of the MS and C rings, the FlgV ring may stabilize interactions between FliG and MotA. There is precedence for auxiliary proteins modulating interactions between the stator and rotor, such as the *Bacillus subtilis* motor clutch proteins EpsE and MotI, which disengage the stator units from the rotor (Blair *et al*, 2008; Subramanian *et al*, 2017).

Alternatively, the FlgV ring may assist the switch complex in controlling rotation direction of the flagellar rotor. In the bacterial chemotaxis system, binding of repellents and attractants to chemoreceptors modulate the autophosphorylation activity of CheA, which is required for phosphorylation of CheY (Jimenez-Pearson *et al*, 2005; Wadhams & Armitage, 2004). Binding of CheY-phosphate to the switch complex causes the rotor to change the direction it rotates from counterclockwise (CCW) to clockwise (CW), resulting in a reorienting of the cell (Karmakar, 2021; Wadhams & Armitage, 2004). The CheY-phosphate-induced directional switch involves a conformational change in FliG that causes the FliG Helix_Torque_ to engage the opposite side of the MotA pentamer (Carroll *et al*, 2020; Chang *et al*, 2020; Santiveri *et al.*, 2020), which results in the MotA pentamer powering the rotation of the rotor in the CW direction. Given the proximity of the FlgV ring to FliG, the FlgV ring may aid in motor reversals that might be important for agar-based motility. Examining the swimming behavior of the *H. pylori* Δ*flgV* mutants and the Δ*flgV* motile variants should allow us to distinguish between these models for FlgV function.

A serendipitous result from our study was the finding that the *H. pylori* G27M motor lacks the FliL_C_ ring (Figure 5K and 5L), which was interesting since *fliL* is reported to be required for motility in *H. pylori* (Tachiyama *et al.*, 2022). A subsequent report by Liu and co-workers, however, indicated FliL is not required for motility in *H. pylori*, but the site where *fliL* is disrupted does impact motility (Liu *et al*, 2022). The authors reported a Δ*fliL* mutation that expresses a truncated FliL protein that lacked FliL_C_ but retained an intact TM domain inhibited motility, whereas motility was enhanced in a Δ*fliL* mutant lacking the entire TM domain (Liu *et al.*, 2022). The report by Liu and co-workers appears to contradict the observation that *H. pylori* G27M has a nonsense mutation in codon 78 of *fliL* and is predicted to express a truncated FliL protein containing the N-terminal cytoplasmic region, intact TM domain, and linker region. The apparent discrepancy may be due to the site of the FliL truncation. In G27M, the FliL truncation occurs at codon 78, whereas in the *H. pylori* Δ*fliL3* mutant generated by Liu and co-workers the truncation is at codon 51 (Liu *et al.*, 2022). Alternatively, the truncated FliL protein expressed in G27M may be turned over rapidly, which would mitigate any inhibitory effect the truncated FliL protein might have on motility.

Regardless of the reason for the apparent discrepancy, it is clear that FliL_C_ is not required for motility in *H. pylori*. FliL_C_ is a member of the SPFH protein family and has been suggested to regulate the activity of the stator units (Tachiyama *et al.*, 2022; Takekawa *et al.*, 2019). We postulate that in the absence of FliL_C_, the unregulated activity of the stator units results in a constitutively high level of torque, which accounts for the enhanced motility of the *H. pylori* G27M strain in soft agar medium.

Although we isolated Δ*flgV* variants that had robust motility in soft agar medium in both the G27M and B128 backgrounds, we were unable to identify specific mutations that were responsible for rescuing motility. It is possible the mutations that suppressed the motility defect of the Δ*flgV* are phase variable and rapidly reverted to the wild-type allele as the Δ*flgV* motile variants were propagated. Only a small fraction of cells within a population would presumably need to maintain the suppressor mutation to generate a large swim halo following inoculation into the soft agar medium. It does not seem coincidental that all three G27M Δ*flgV* motile variants had a mutation in *flaG-*2, which resulted in many of the cells having significantly longer flagellar filaments (Figure 3B and 3C, Table 2). Although the *flaG*-*2313* allele by itself failed to rescue motility in the G27M Δ*flgV* mutant, the *flaG*-2 mutation may contribute to rescuing motility in the G27M Δ*flgV* motile variants since stator units were not apparent in the motor of the G27M Δ*flgV* mutant (Figure 5K). In contrast, stator units were visible in the motors of the G27M strain expressing the FlgV-YFP fusion protein (Figure 5J), G27M Δ*flgV* mutant complemented with *flgV* (Figure S3), and B128 Δ*flgV* mutant (Figure 5I). Taken together, these observations suggest that in the absence of FlgV and FliL_C_ the stator units are not retained stably within the motor. In the *E. coli* motor, stator units within the motor exchange readily with a pool of unbound stators in the membrane, and the number of stators that engage the rotor varies according to the load on the motor (Lele *et al.*, 2013). The abnormally long flagellar filaments of the G27M Δ*flgV* motile variants may therefore help to rescue motility by increasing the load to facilitate recruitment of stators to the *H. pylori* motor.

## MATERIALS AND METHODS

### Bacterial strains and growth conditions

*E. coli* NEB TURBO was used for cloning and plasmid construction and was grown in LB liquid or agar medium supplemented with ampicillin (100 μg/ml) or kanamycin (30 μg/ml) when appropriate. *H. pylori* strains were grown microaerobically under an atmosphere consisting of 10% CO_2_, 4-6% O_2_ and 84-86% N_2_ at 37°C on tryptic soy agar supplemented with 5% horse serum (TSA-HS) or Columbia agar supplemented with 5% sheep blood. Liquid cultures of *H. pylori* were grown in Brain Heart Infusion (BHI) medium supplemented with 5% heat-inactivated horse serum with shaking in serum bottles under an atmosphere consisting of 5% CO_2_, 10% H_2_, 10% O_2_ and 75% N_2_. *H. pylori* growth medium was supplemented with kanamycin (30 μg/ml) or sucrose (5%) where appropriate.

### Construction of *H. pylori* Δ*flgV* mutants

All primers used for PCR in the construction of *H. pylori* mutants are listed in Table S6. Plasmids and *H. pylori* stains used in the study are listed in Table 4. Genomic DNA from *H. pylori* G27 was purified using the Wizard genomic DNA purification kit (Promega, Madison, WI, USA) and used as the PCR template to construct the suicide vectors used to generated deletion mutants. PCR was carried out using Phusion DNA polymerase (New England Biolabs, Ipswich, MA, USA) and the resulting amplicons were incubated with *Taq* polymerase (Promega) at 72° C for 10 min to add 3’-A overhangs for T/A cloning with pGEM-T Easy plasmid (Promega). A 610 bp DNA fragment upstream of *flgV* was amplified using PCR primers 85 and 86, and a 600 bp downstream DNA fragment was amplified using PCR primers 87 and 88. PCR SOEing was used to join the upstream and downstream regions, and the resulting amplicon was cloned into pGEM-T Easy to generate plasmid pKHG25. Plasmid pKHG25 was cut at unique XhoI and NheI restriction sites and ligated with a kan^R^*-sacB* cassette (Chu *et al*, 2019) to generate plasmid pKHG26. The suicide vector pKHG26 was introduced into *H. pylori* G27M or *H. pylori* B128 by natural transformation. Transformants in which the kan^R^*-sacB* cassette on the plasmid had replaced the chromosomal copy of *flgV* by allelic exchange were selected on TSA-HS supplemented with kanamycin. Replacement of the kan^R^*-sacB* cassette with *flgV* in kanamycin-resistant transformants was confirmed by PCR, and the resulting strains were designated KHG40 (B128 mutant) and KHG45 (G27M mutant). The suicide vector pKHG25 was introduced into strains KHG40 and KHG45 by natural transformation, and sucrose-resistant transformants in which the unmarked *flgV* deletion carried in plasmid pKHG25 had replaced the kan^R^*-sacB* cassette in the chromosomal copy of *flgV* by allelic exchange were isolated by plating onto TSA-HS supplemented with 5% sucrose. Sucrose-resistant colonies were screened for sensitivity to kanamycin, and the deletion of *flgV* in a kanamycin-sensitive isolate was confirmed by PCR and DNA sequencing (Eton Bioscience, Research Triangle Park, NC, USA) of the resulting amplicon. The resulting G27M and B128 Δ*flgV* mutants were designated KHG48 and KHG62, respectively.

**Table 4.**
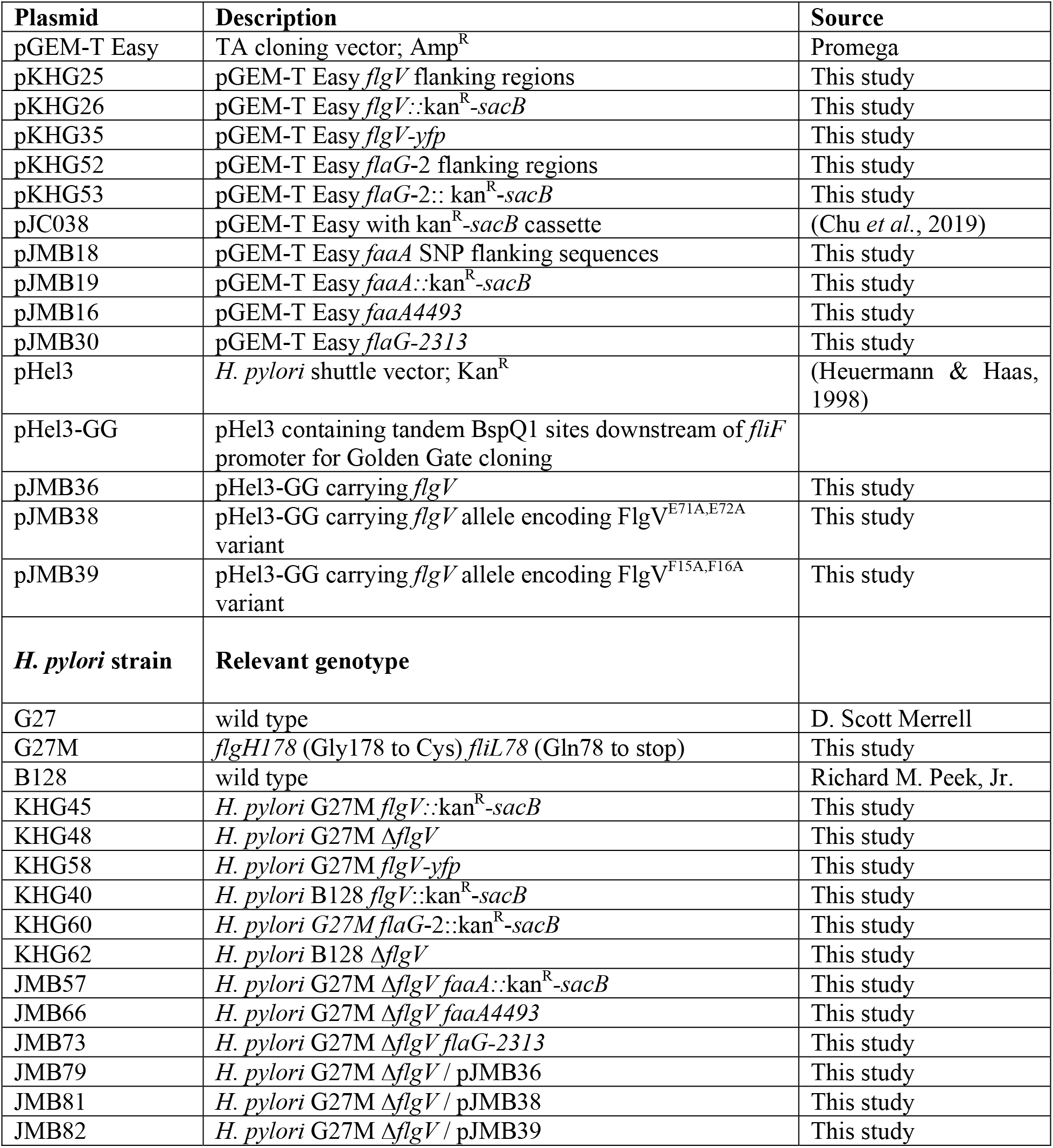
Plasmids and *H. pylori* strains used in this study.

| Plasmid | Description | Source |
| --- | --- | --- |
| pGEM-T Easy | TA cloning vector; Amp<sup>R</sup> | Promega |
| pKHG25 | pGEM-T Easy *flgV* flanking regions | This study |
| pKHG26 | pGEM-T Easy *flgV::kan<sup>R</sup>-sacB* | This study |
| pKHG35 | pGEM-T Easy *flgV-yfp* | This study |
| pKHG52 | pGEM-T Easy *flaG-2* flanking regions | This study |
| pKHG53 | pGEM-T Easy *flaG-2::kan<sup>R</sup>-sacB* | This study |
| pJC038 | pGEM-T Easy with kan<sup>R</sup>-*sacB* cassette | (Chu *et al.*, 2019) |
| pJMB18 | pGEM-T Easy *faaA* SNP flanking sequences | This study |
| pJMB19 | pGEM-T Easy *faaA::kan<sup>R</sup>-sacB* | This study |
| pJMB16 | pGEM-T Easy *faaA4493* | This study |
| pJMB30 | pGEM-T Easy *flaG-2313* | This study |
| pHel3 | *H. pylori* shuttle vector; Kan<sup>R</sup> | (Heuermann & Haas, 1998) |
| pHel3-GG | pHel3 containing tandem BspQ1 sites downstream of *fliF* promoter for Golden Gate cloning |  |
| pJMB36 | pHel3-GG carrying *flgV* | This study |
| pJMB38 | pHel3-GG carrying *flgV* allele encoding FlgV<sup>E71A,E72A</sup> variant | This study |
| pJMB39 | pHel3-GG carrying *flgV* allele encoding FlgV<sup>F15A,F16A</sup> variant | This study |
| *H. pylori* strain | Relevant genotype |  |
| G27 | wild type | D. Scott Merrell |
| G27M | *flgH178* (Gly178 to Cys) *fliL78* (Gln78 to stop) | This study |
| B128 | wild type | Richard M. Peek, Jr. |
| KHG45 | *H. pylori* G27M *flgV::kan<sup>R</sup>-sacB* | This study |
| KHG48 | *H. pylori* G27M Δ*flgV* | This study |
| KHG58 | *H. pylori* G27M *flgV-yfp* | This study |
| KHG40 | *H. pylori* B128 *flgV::kan<sup>R</sup>-sacB* | This study |
| KHG60 | *H. pylori* G27M *flaG-2::kan<sup>R</sup>-sacB* | This study |
| KHG62 | *H. pylori* B128 Δ*flgV* | This study |
| JMB57 | *H. pylori* G27M Δ*flgV faaA::kan<sup>R</sup>-sacB* | This study |
| JMB66 | *H. pylori* G27M Δ*flgV faaA4493* | This study |
| JMB73 | *H. pylori* G27M Δ*flgV flaG-2313* | This study |
| JMB79 | *H. pylori* G27M Δ*flgV* / pJMB36 | This study |
| JMB81 | *H. pylori* G27M Δ*flgV* / pJMB38 | This study |
| JMB82 | *H. pylori* G27M Δ*flgV* / pJMB39 | This study |

### Complementation of *flgV* mutation

To facilitate complementation assays in *H. pylori*, we modified the shuttle vector pHel3 (Heuermann & Haas, 1998) for Golden Gate assembly using tandem sites for the Type IIS restriction enzyme BspQ1. The modified plasmid, which we designated pHel3-GG, contains the *fliF* promoter and the *ureA* ribosome-binding site upstream of the tandem BspQ1 sites where the cloned gene is introduced. Primers 151 and 152 were used to amplify *flgV* from *H. pylori* G27 genomic DNA. The primers introduced BspQ1 sites immediately upstream and downstream of the start and stop codons of *flgV*, respectively. The resulting amplicon and pHel3-GG were digested together with BspQ1 (New England Biolabs) for 1 h at 50°C, after which the amplicon and vector were ligated using Fast-Link DNA ligase (Biosearch Technologies, Hoddesdon, UK) for 30 min at room temperature. The reaction mix was incubated with additional BspQ1 for 1 h at 50°C, then used for transformation of *E. coli*. A plasmid containing the expected insert was identified following restriction enzyme digestion and the identity of the insertion was confirmed by DNA sequencing (Eton Biosciences). The resulting plasmid, pJMB36, was introduced into KHG48 by natural transformation to generate strain JMB79.

### Site-directed mutagenesis of *flgV*

Specific mutations were introduced into *flgV* carried on plasmid pJMB36 using the primer pairs 153 and 154, 155 and 156, or 157 and 158 together with the Q5 Site-Directed Mutagenesis Kit (New England Biolabs) as per the manufacturer’s instructions. The resulting plasmids, pJMB39 and pJMB38, were individually introduced into strain KHG48 by natural transformation to generate strains JMB81 and JMB82.

### Construction of *H. pylori* strain expressing FlgV-YFP fusion

PCR primers 85 and 74 amplified *flgV* and an upstream flanking region totaling 877 bp, omitting the stop codon of *flgV*. Primers 75 and 76 were used to amplify *yfp* and added a sequence encoding a flexible linker (Gly-Ser-Ala-Gly-Ser-Gly) that joined *flgV* and *yfp*. Primers 77 and 78 amplified a 512 bp downstream sequence of *flgV*. PCR SOEing was used to join the all three fragments, and the resulting amplicon was cloned into pGEM-T Easy to generate the suicide vector pKHG35. Plasmid pKHG35 was introduced to strain KHG45 by natural transformation, and sucrose-resistant transformants in which the kan^R^*-sacB* cassette in the chromosomal copy of *flgV* had been replaced with the *flgV*-*yfp* fusion by allelic exchange were isolated on TSA-HS supplements with kanamycin. The presence of the *flgV-yfp* fusion in the *flgV* locus was confirmed by PCR, and the strain was designated KHG58.

### Motility assay

Motility of *H. pylori* strains in a soft agar medium consisting of Mueller-Hinton broth, 10% heat-inactivated horse serum, 20 mM 2-(N-morpholino)ethanesulfonic acid (MES; pH 6.0), and 0.4% Noble agar was done as described (Chu *et al.*, 2019). Diameters of the swim halos emanating from the point of inoculation in the soft agar medium were measured 7 d post-inoculation. A minimum of three replicates were done for each strain. Mean values for the of the swim halo diameters were calculated and a two-sample *t* test was used to determine statistical significance.

### Isolation of motile variants of Δ*flgV* mutant

Enrichment of motile variants of the *H. pylori* G27M Δ*flgV* mutant was accomplished by six rounds of inoculating the mutant in soft agar medium, allowing the bacteria to migrate from the point of inoculation, picking cells from the edge of the swim halo, and inoculating fresh soft agar medium. Following the last iteration, cells from the edge of the swim halo were streaked onto TSA-HS to obtain single colonies. Three independent enrichments were done and a single motile isolate from each enrichment was saved and characterized.

### Introduction of *faaA* and *flaG*-2 alleles into the *H. pylori* G27M Δ*flgV* mutant

PCR primer pair 147 and 148 amplified 526 bp within *faaA*, and primer pair 149 and 150 amplified 534 bp within *faaA*. The two amplicons were joined by PCR SOEing, and the resulting fragment was cloned into pGEM-T Easy to generate plasmid pJMB18. The kan^R^-*sacB* cassette was introduced into plasmid pJMB18 at unique XhoI and NheI restriction sites to generate plasmid pJMB19, which was introduced into strain KHG48 by natural transformation. Transformants in which the kan^R^*-sacB* cassette on the plasmid had been incorporated into the chromosomal copy of *faaA* by allelic exchange were selected on TSA-HS supplemented with kanamycin. The presence of the kan^R^*-sacB* cassette within *faaA* was confirmed by PCR, and the resulting strain was designated JMB57. Primers 147 and 150 were used to amplify 1658 bp including the *faaA* SNP identified in MV2. The amplicon was cloned into pGEM-T Easy to generate plasmid pJMB16, which was introduced into strain JMB57 by natural transformation. Sucrose-resistant transformants in which the *faaA* allele carried in plasmid pJMB16 had replaced the kan^R^*-sacB* cassette in the chromosomal copy of *faaA* by allelic exchange were isolated by plating onto TSA-HS supplemented with 5% sucrose. Sucrose-resistant colonies were screened for sensitivity to kanamycin, and introduction of the mutant *faaA* allele in the chromosome of a kanamycin-sensitive isolate was confirmed by PCR and DNA sequencing (Eton Bioscience). The resulting strain was designated JMB66.

The *flaG*-*2313* allele was introduced into *H. pylori* G27M Δ*flgV* as follows. A 592 bp DNA sequence upstream of *flaG*-2 was amplified using PCR primers 107 and 108, and a 581 bp DNA sequence downstream of *flaG*-2 was amplified using PCR primers 109 and 110. PCR SOEing was used to join the upstream and downstream regions, and the resulting amplicon was cloned into pGEM-T Easy to generate plasmid pKHG52. The kan^R^*-sacB* cassette was cloned into plasmid pKHG52 at unique XhoI and NheI restriction sites to generate the suicide vector pKHG53, which was introduced into strain KHG48 by natural transformation. Presence of the kan^R^*-sacB* cassette in kanamycin-resistant transformants was confirmed by PCR, and the resulting strain was designated KHG60. Primers 107 and 110 were used to amplify 438 bp that include the *flaG*-2 SNP identified in MV1 and MV3 (i.e., *flaG*-*2313*). The resulting amplicon was cloned into pGEM-T Easy to generate the suicide plasmid pJMB30, which was introduced into KHG60 by natural transformation. Sucrose-resistant transformants in which the *flaG-*2 allele carried on plasmid pJMB30 had replaced the kan^R^*-sacB* cassette in the chromosomal copy of *flaG-*2 by allelic exchange were isolated by plating onto TSA-HS supplemented with 5% sucrose. Sucrose-resistant colonies were screened for sensitivity to kanamycin, and introduction of the *flaG*-*2313* allele in the chromosome of a kanamycin-sensitive isolate was confirmed by PCR and DNA sequencing (Eton Bioscience). The resulting strain was designated JMB73.

### Transmission electron microscopy

*H. pylori* cultures were grown to late-log phase (A_600_ = ~1.0) in BHI supplemented with 10% heat-inactivated horse serum. Cells from the cultures were collected by centrifugation, fixed with formaldehyde and glutaraldehyde, and negatively stained with uranyl acetate as described (Chu *et al.*, 2019). Cells were visualized using a JEOL JEM 1011 transmission electron microscope operated at 80 kV. Flagella lengths were measured from TEM images using Fiji software (Schindelin *et al*, 2012), and length profiles were compared using a Mann-Whitney U test.

### Genome sequencing and analysis

Genomic libraries, prepared with the Illumina iTruSeq adaptor kit from 500 ng of gDNA from various *H. pylori* strains, were sequenced at either the University of Georgia Genomics Facility or Azenta Life Sciences (Chelmsford, MA, USA) by Illumina sequencing. Alternatively, gDNA was provided to the Seqcenter (Pittsburgh, PA, USA), which prepared genomic libraries and sequenced the libraries by Illumina sequencing. Sequence analysis was performed using the *breseq* computational pipeline (Deatherage & Barrick, 2014). A reference genome was constructed by mapping reads on an NCBI genome for *H. pylori* B128 (Accession: NC_011333.1). Reads were mapped to the reference genome, and a minimum variant frequency of 0.8 was used to call variants and identify SNPs. Variants and SNPs in the strains were compared manually to identify ones that were present only in the suppressor strains.

### Cryo-ET sample preparation

*H. pylori* strains were grown on Columbia agar plates supplemented with 5% horse red blood cells at 37 °C in 10% CO_2_ conditions. For the complemented strains, the medium was supplemented with 30 μg/mL of kanamycin to maintain the shuttle vector. Bacteria from the agar medium were resuspended in phosphate buffer saline (PBS) and mixed with 10 nm of BSA gold tracers (Aurion, Wageningen, NL). The mixtures were deposited on freshly grow-discharged cryo-EM grids (Quantifoil R2/1, Cu 200, Ted Pella, Inc., Redding, CA, USA), and then frozen into liquid ethane by using a manual plunger.

### CryoET data collection and processing

Frozen-hydrated specimens were visualized in a 300kV Titan Krios electron microscope (Thermo Fisher Scientific, Waltham, MA, USA) equipped with a K3 summit direct detection camera and a BioQuantum energy filter (Gatan, Pleasanton, CA, USA). Tilt series were acquired at 42,000x magnification (corresponding to a pixel size of 2.148 Å at the specimen level) by using SerialEM (Mastronarde, 2005) and FastTomo script (Xu *et al*. 2021) based on a dose-symmetric scheme at defocus ~4.5 μm. The stage was tilted from −48° to +48° at 3° increments. The total accumulative dose for each tilt series was ~60 e^−^/Å^2^. Motioncorr2 was used for drift correction (Zheng *et al*, 2017). Gold tracer beads were tracked to align all image stacks using IMOD (Kremer *et al*, 1996). Gctf was used for all aligned stacks to estimate defocus (Zhang 2016), and then the ctfphaseflip function in IMOD was used for contrast transfer function (CTF) correction.

### Subtomogram averaging

Tomo3D (Agulleiro and Fernandez, 2015) was used for 3D reconstruction, and a total of 268, 420, 224, 332, and 214 tomograms were reconstructed from the aligned tilt series of the wild-type, *ΔflgV* with *ΔfliL* (G27M), *ΔflgV* (B128), FlgV-YFP, and complemented strains, respectively. Binned 6 × 6 × 6 tomograms using the simultaneous iterative reconstructive technique (SIRT) were used to select 824, 2523, 654, 1255, and 659 flagellar motors at the bacterial pole from the wild-type, *ΔflgV* with *ΔfliL* (G27M), *ΔflgV* (B128), FlgV-YFP, and complemented strains respectively. After picking particles, tomograms with weighted back projection (WBP) were used for the initial subtomogram average structures using i3 suite (Winkler, 2007; Winkler *et al*, 2009). Binned 4 × 4 × 4 subtomograms were used to refine the intact motor structures. Classification was used to remove non-flagellated motors.

## ACKNOWLEDGEMENTS

We thank Winsen Wijaya for his technical assistance. We thank Jennifer Aronson for critical reading of the manuscript. This work was supported by NIH grants AI140444 and AI146907 to T.R.H. and AI087946 and AI132818 to J.L.

